# Neuromuscular Basis of Kinematic Adaptations During Bidirectional Walking

**DOI:** 10.64898/2026.02.11.705376

**Authors:** Helia Mojtabavi, Atra Ajdari, Sebastian Rueda-Parra, Darren E. Gemoets, Jonathan R. Wolpaw, Russell L. Hardesty

**Affiliations:** National Center for Adaptive Neurotechnologies (NCAN), Albany, NY; Department of Veterans Affairs, Samuel S. Stratton Medical Center, Albany, NY

**Keywords:** split-belt treadmill, locomotor adaptation, bidirectional walking, muscle activation, EMG, motor learning, neuromuscular control

## Abstract

1

**Background:** Human locomotion is a highly adaptive motor skill that adjusts to new environmental demands through learning. Split-belt treadmill paradigms have advanced our understanding of gait adaptation. Most studies have examined gait when the belts move at different speeds in the same direction. We are studying muscle activation patterns during an asymmetric gait, when the treadmill belts move at equal speed in opposite directions, i.e., bidirectional walking (BDW).

**Methods:** Twelve healthy volunteers performed a single session on a split-belt treadmill. We simultaneously collected ground reaction forces via treadmill force plates, joint kinematics via motion capture, and surface electromyography (EMG) from bilateral soleus (SOL) and tibialis anterior (TA) muscles. Participants started with 2 min of forward walking (FW), followed with four 5-min blocks of BDW separated by 1-min standing rest intervals, and finished the session with 2 min of FW (washout).

**Results:** All participants successfully completed the protocol. We analyzed EMG signals for temporal activation patterns (rhythm generation) and amplitude characteristics (pattern formation). EMG recordings revealed antiphasic activation of SOL and TA muscles bilaterally throughout all trials. During BDW, the backward-moving leg’s TA showed prolonged activation patterns that persisted during washout FW, suggesting retention of adaptive changes. Burst-to-cycle duration ratios showed transient changes during early adaptation but remained relatively stable across conditions, demonstrating robust rhythm generation despite adaptive modulation of activation patterns during BDW.

**Discussion:** These findings demonstrate that BDW induces asymmetric adjustments in muscle activation patterns. Rhythm generation (timing) did not significantly differ between BDW and FW. However, we did observe changes in pattern formation (i.e., EMG profiles) during FW pre- and post-BDW training. Burst-to-cycle duration ratios, as a measure of rhythm generation, showed changes during early adaptation, particularly the increase in right SOL and right TA during block 1, though these changes did not reach statistical significance and largely returned to baseline during washout. The underlying pattern formation structure, was maintained across all conditions, with selective amplitude modulations rather than fundamental reorganization of activation patterns. The substantial temporal adjustments in the backward-moving leg’s SOL and phase shifts in TA provide the neuromuscular mechanism driving the bilateral step-length reduction, altered inter-limb phasing, and asymmetric double stance timing. These results extend our understanding of locomotor control by suggesting how the central nervous system (CNS) dynamically recalibrates muscle timing and amplitude to maintain satisfactory locomotion under new environmental demands.

## 2 Introduction

Human locomotion is a complex and highly adaptive behavior. Continuous adjustments are necessary during everyday actions like turning, running, or walking on challenging surfaces to ensure that function well in the changing environments. This responsive flexibility is managed by complex neuromuscular coordination. The hierarchy of human locomotion control is often described as a tri-partite system comprised of spinal central pattern generator (CPG), brainstem, and cortical structures [1]. In the CPG, locomotor control is organized hierarchically into a rhythm generator responsible for the temporal structure and timing of muscle activation and a pattern formation layer that determines the amplitude of muscle output. Somatosensory feedback from the environment, along with descending commands from the brainstem and higher cortical regions, modulate these rhythm and pattern mechanisms to adjust human gait and meet current environmental demands [2, 3, 4, 1].

The introduction of split-belt treadmill paradigms made studying the specifics of gait adjustments in diverse conditions possible [5]. These studies have focused predominantly on spatiotemporal parameters of gait (e.g., step length and symmetry, and center of pressure, and phase of oscillation), with limited attention to muscle activation properties [6, 7, 8, 9, 10, 11]. Studying temporal activation patterns (rhythm generation) and amplitude characteristics (pattern formation) via electromyography (EMG) is essential because it provides mechanistic insight into the underlying neural circuits and muscle synergies that govern human locomotion. Furthermore, most existing studies focus on how gait adapts to differences in speed in one direction, leaving adaptation to opposite walking in opposite directions (i.e., bidirectional walking (BDW), also referred to hybrid locomotion) understudied.

BDW poses a unique challenge to the locomotor control system and is commonly observed in day-to-day life when animals or people pivot or perform a sharp turn [12]. Studies in decerebrated or spinalized cats demonstrate that spinal circuits can generate BDW without forebrain control [13, 12]. Crucially, while adapting to bidirectional coordination, the central networks showed no significant differences in cycle durations between the forward and backward-stepping limbs, suggesting a common rhythm-generating mechanism within the spinal circuits [13, 12].

However, studying adaptation in healthy adult humans using the split-belt treadmill reveals direction-specific results. Choi and Bastian [14] found that motor adaptations learned during forward walking (FW) do not transfer to backward walking (BW), and adaptations learned during BW do not transfer to FW. These findings suggest that FW and BW in humans are controlled by separate functional networks [14]. While these results highlight directional independence, subsequent research on humans suggests that the motor system still maintains strong bilateral interactions between the limbs during adaptation, as proximal muscle activity is influenced by the speed of the contralateral belt [15]. In contrast, studies in feline models have suggested a more shared spinal network for both walking directions, as they exhibit similar muscle synergies and spatiotemporal strategies for speed modulation in both forward and backward walking [16].

Here, we investigate the adaptive changes in muscle activity in response to bidirectional walking. Twelve healthy participants performed a single session on an instrumented split-belt treadmill, in which the belts were moving at equal speeds but in opposing directions. We simultaneously collected ground reaction forces (kinetic data) and joint trajectories (kinematic data) via passive motion capture markers, along with surface EMG from the antagonistic muscles, soleus (SOL) and tibialis anterior (TA), bilaterally. In our initial article [17], we reported on the immediate changes of spatiotemporal gait characteristics, such as step length asymmetry, to BDW. Here we test the hypothesis that bilateral muscle activation patterns adapt to accommodate the asymmetric demands of BDW by: (1) quantifying the temporal activation (timing/rhythm generation) and intensity (amplitude/pattern formation) of EMG signals; and (2) correlating the EMG results with the kinematic results.

## 3 Methods

### 3.1 Ethical approval

All experimental procedures were approved by the Institutional Review Board (IRB) of the Stratton VA Medical Center and conducted in accordance with the *Declaration of Helsinki* (Protocol #:1584762). All participants provided written informed consent prior to participation.

### 3.2 Participants

We recruited twelve healthy volunteers for this study. Inclusion criteria required that participants had no neurological deficits, were able to walk independently on a treadmill for 30 min, and had no prior experience with split-belt treadmill walking. We collected basic demographic information including age, weight, height, and physical activity levels. We assessed foot dominancy using the Waterloo Footedness Questionnaire. Because all participants demonstrated either mixed or right-foot dominance, we assigned the right leg to move in the reverse direction for all participants.

### 3.3 Experimental Paradigm

Participants walked on a split-belt treadmill (Bertec Version 2.0; Columbus, OH) while wearing a non-weight bearing safety harness attached to the treadmill frame. First, the participants were presented five different speeds at ±0.2 m/s increments on a tied-belt treadmill to determine their preferred walking speed (PWS). We then kept the belt’s speed at 80% of PWS throughout the session. Participants started the experiment with 2 min of FW. Then we introduced them to the BDW by gradually (over 1 min) decreasing the right belt’s speed to zero and increasing the speed in the reverse direction to match that of the left leg in absolute value. Following this familiarization period, participants walked for four 5-min blocks of BDW separated by 1-min standing rest periods, and finished the session with another 2-min block of FW. Figure 1 shows the walking protocol.

**Figure 1:**
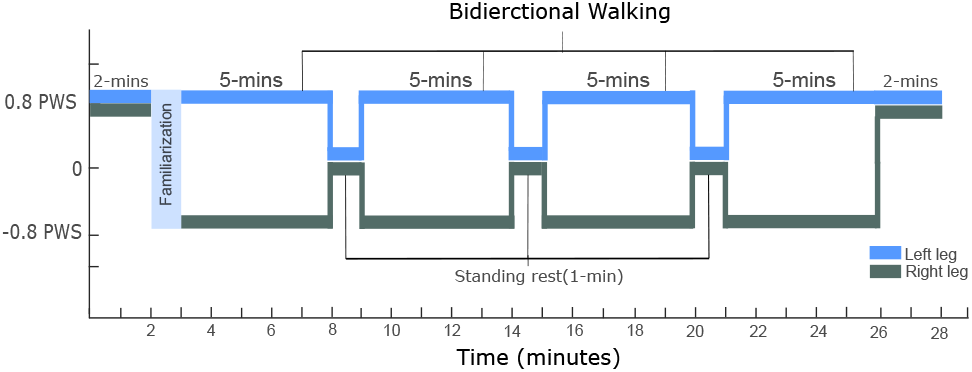
Walking Paradigm. The session began with 2 min of FW to capture baseline measurements. Participants then completed 20 min of bidirectional walking training, divided into four 5-min blocks by 1-minute standing rest periods. All walking was performed at 80% of the participant’s preferred speed. The session concluded with 2 min of FW to measure washout effects.

### 3.4 Data Acquisition

We recorded muscle activity with surface EMG from SOL and TA bilaterally with Ag-AgCl electrodes (Vermed NeuroPlus Cloth EM-1041-0060). For the TA muscle, two electrodes were placed with 2.5-3 cm center-to-center separation along a line parallel to the muscle belly 1 cm lateral to the tibia, and 1/3 of the distance from the lateral epicondyle of the femur and the lateral malleolus. For SOL muscle, two electrodes were placed on the midline 2 cm the lower edge of gastrocnemius. EMG was amplified (gain 500) and band-pass filtered (10–1000 Hz) (AMT-8 EMG amplifier, Bortec Biomedical Ltd., Calgary, Alberta, Canada), and then digitized (3200 Hz).

Ground reaction forces (GRF) and moments were recorded concurrently in three dimensions with the two force plates of the treadmill at a frequency of 2000 Hz. GRF, moments, and EMG were stored in the same computer with our Qualisys Track Manager (QTM) software and its data acquisition board.

### 3.5 Data processing

#### EMG parameters

First, we applied in sequence a fourth-order Butterworth 20-Hz high-pass filter to remove movement artifacts and baseline drift [18] a 60-Hz notch filter to eliminate power line interference, and a denoising procedure to reduce residual noise while preserving the underlying muscle activation patterns. We then mean-centered each signal by subtracting its trial-specific mean amplitude and performed full-wave rectification by computing the absolute value of the voltage signal. To extract the EMG amplitude envelope, we applied a fourth-order Butterworth low-pass filter with a cutoff frequency of 30 Hz to the rectified signal (Figure 2). This smoothing procedure preserved movement-related amplitude modulations while attenuating residual high-frequency noise components.

**Figure 2:**
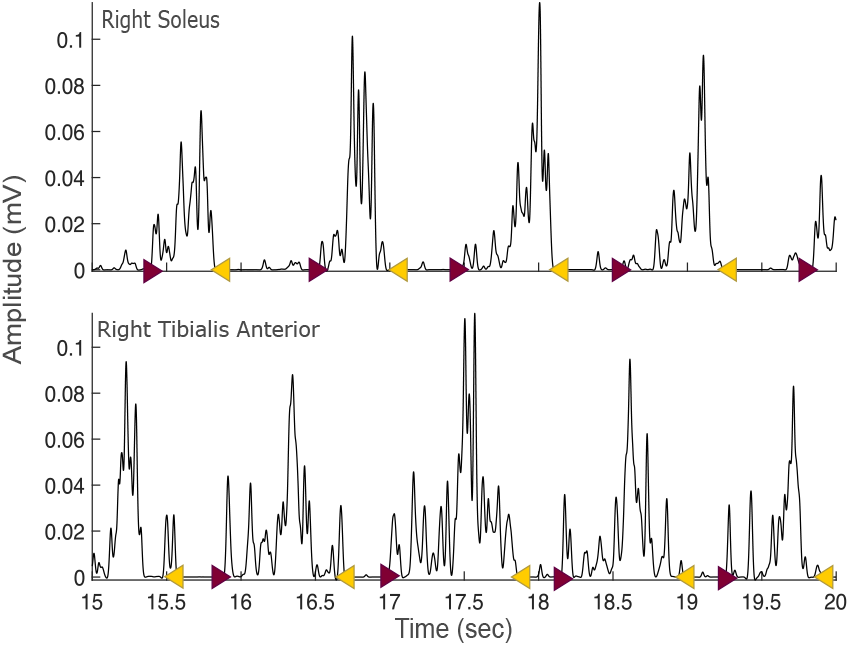
Representative EMG burst timing during bidirectional walking. EMG signals from right SOL and TA during a segment of bidirectional walking. Purple arrows indicate burst onsets and yellow arrows indicate burst offsets, which were manually identified from the rectified EMG signals.

##### Cycle duration

To calculate cycle duration, we detected step onsets and offsets and then applied a second-order 20-Hz low-pass Butterworth filter to the force plate measurements in the vertical (Z) direction to reject ambient electrical noise and other high frequency interference. We thresholded the GRF signal to detect onsets and offsets for each leg. We then calculated cycle duration as the time difference between successive right foot onsets.

##### Amplitude

Time-normalized EMG envelopes were compared between the Baseline and Washout FW and Block 14 of BDW at the group level using a cluster-based permutation test [19, 20, 21]), a widely used approach in electroencephalography (EEG), EMG, and fMRI analyses for assessing differences in high-dimensional, and time-series data.

Amplitude analysis comparison followed three main steps:

##### Step 1. Permutation of condition labels and generation of surrogate differences

Condition labels (Baseline and Washout) were randomly permuted within each participant 1000 times. For every permutation, condition-specific EMG envelopes were averaged across participants and subtracted (i.e. mean permuted Washout FW *–* mean permuted Baseline FW), producing 1000 surrogate difference time series representing the assumption of chance differences.

##### Step 2. Construction of the null distribution of maximum cluster sizes

Each permuted difference time series was converted to z-scores using the null distribution’s mean and standard deviation at each time point. These z-score time series were thresholded at |*z*| *>* 1.96 (corresponding to *p* = 0.05, two-tailed), and temporally contiguous supra-threshold points were grouped into clusters. The maximum cluster size from each permutation yielded a null distribution of maximum cluster sizes. This approach controls the family-wise error rate [19, 20].

##### Step 3. Comparison of observed data to the null distribution

The same procedure was applied to the observed (non-permuted) data. First, EMG envelopes were averaged across participants and subtracted to obtain the observed difference time series (No swapping of labels this time). This time series was z-score transformed using the null distribution parameters derived in Step 2 and thresholded at |*z*| *>* 1.96. Contiguous supra-threshold time points were grouped into clusters. Observed clusters exceeding the 95th percentile of the null distribution were considered statistically significant. Following [21], statistical inference was made at the cluster level, not at individual time points.

#### Statistical analysis

Statistical analyses were performed in Python (version 3.12.4) using the SciPy (version 1.13.1) and StatsModels (version 0.14.2) libraries [22, 23, 24]. For each participant, mean burst-to-cycle duration ratios and TA/SOL slope ratios were estimated for each condition using simple linear regression. Paired t-tests compared baseline FW, each BDW block, and washout to assess condition-dependent changes in muscle activation timing and coordination. Median values and 95% confidence intervals were computed using bootstrap resampling (10,000 iterations). Results were visualized with box-plots and scatter plots to illustrate adaptation, de-adaptation, and inter-limb coordination dynamics across walking conditions.

## 4 Results

All 12 participants successfully completed the study (7 right-foot dominant, 5 mixed dominance, 0 left-foot dominant, as assessed by the Waterloo Footedness Questionnaire). Participants had a mean (±SD) age of 42.3 ± 12.3 years, height of 1.71 ± 0.1 m, weight of 71.5 ± 18.4 kg, and preferred forward walking speed of 0.896 ± 0.2 m/s.

### 4.1 EMG activation pattern during forward and Bidirectional walking

To determine whether BDW would induce changes in muscle activation amplitude (pattern formation) during adaptation and whether these changes would cause aftereffects that persist during FW washout, we examined EMG amplitude envelopes across all conditions. We time-normalized the processed EMG envelopes relative to gait cycle to account for inter-trial variability in movement duration and normalized amplitude values to each participant’s peak EMG amplitude across all trials (Figure 3). This allowed us to compare relative muscle activation levels both within and between participants.

**Figure 3:**
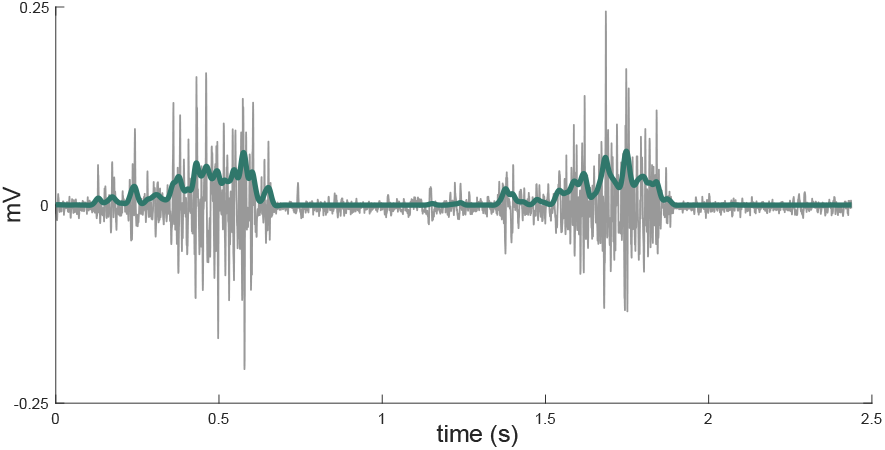
EMG signal processing. Example of raw EMG signal (gray) and its processed envelope (green).

**Figure 4:**
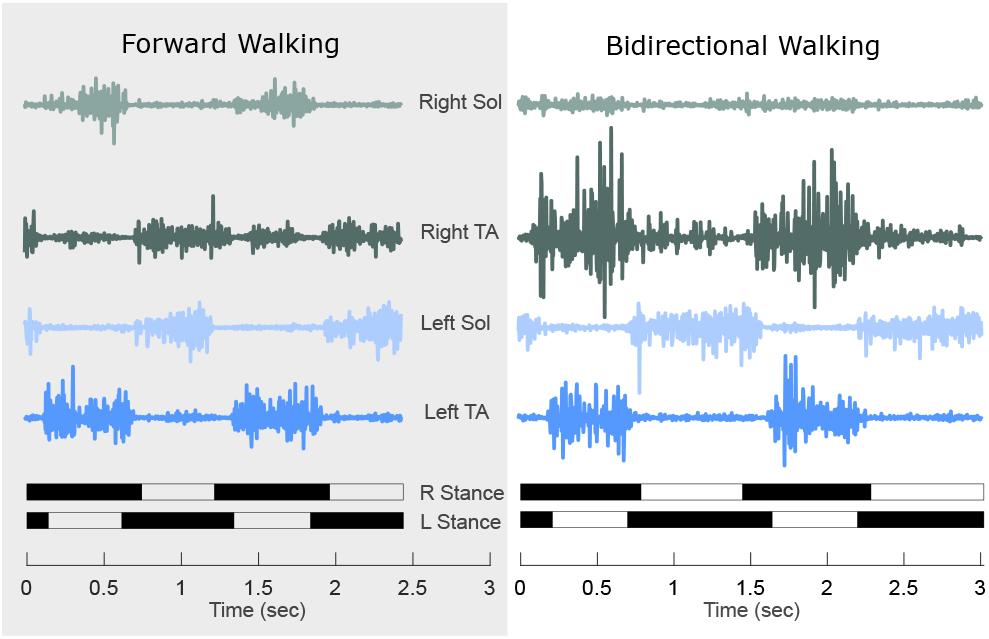
Representative EMG activity during forward and bidirectional walking. Raw EMG signals from the SOL and TA muscles of both legs during forward walking (left panel) and bidirectional walking (right panel) (from a representative participant). Black and white bars at the bottom indicate right (R) and left (L) stance phases. Data shown are from a representative 3-second segment from one of the participants.

During FW, the right SOL showed activation during mid to late stance and push-off at both baseline and washout, with a statistically significant difference between baseline (black line) and washout (gray line) observed during early stance phase (p = 0.027) in the highlighted yellow region (Figure 5 A, first panel). The right TA displayed a bimodal activation pattern with a brief burst following right heel strike during early stance and a larger burst in late swing preceding the next heel strike at both baseline and washout, with no significant differences detected (Figure 5 A, second panel). Left SOL showed activation during mid-to-late stance at baseline and washout (Figure 5 A, third panel), and the left TA displayed a bimodal activation pattern at baseline and washout (Figure 5 A, fourth panel). No significant differences were detected in the right TA, left SOL, or left TA during FW (Figure 5, Row A second to fourth panel).

**Figure 5:**
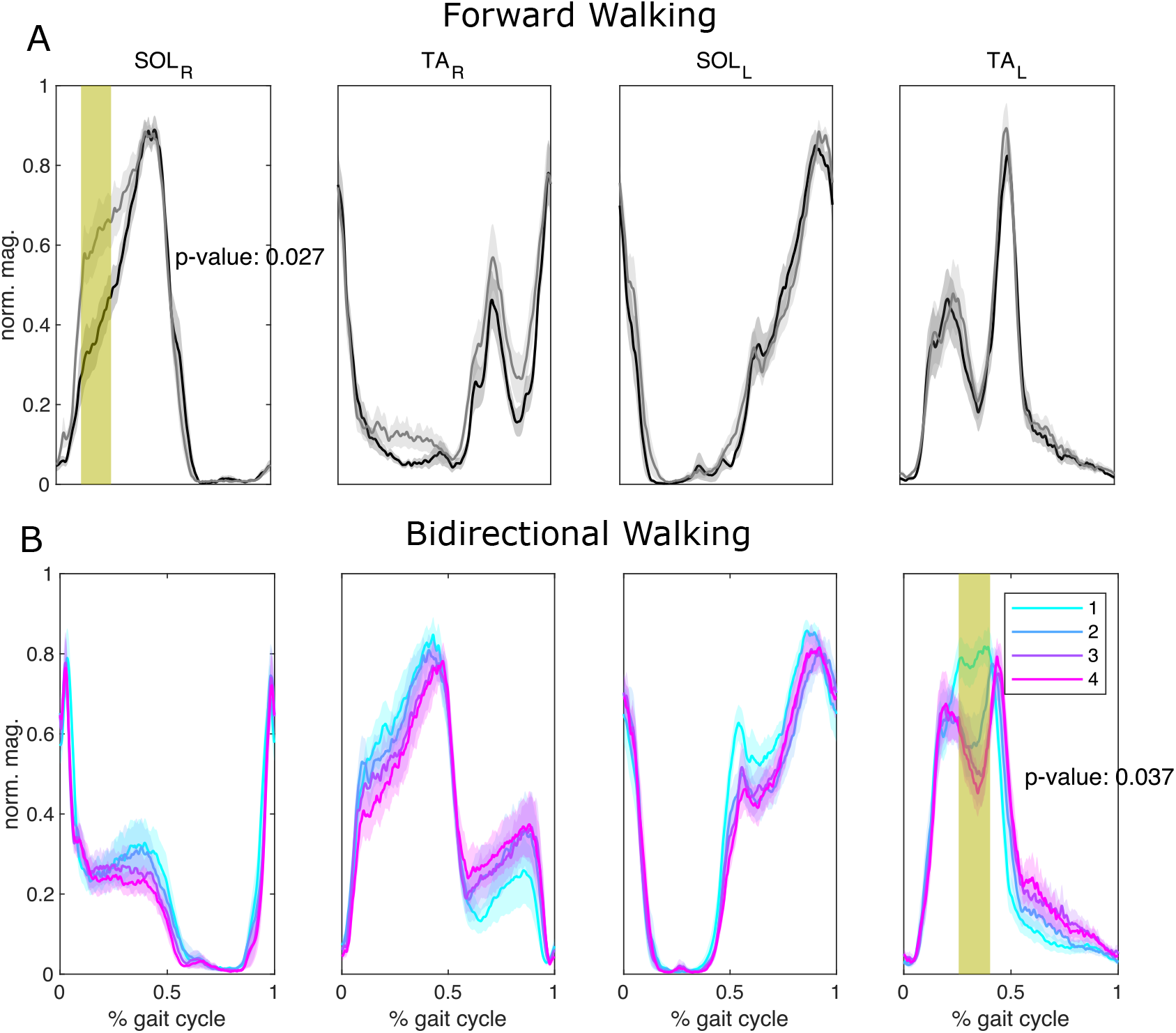
EMG amplitude differences between walking blocks. Normalized EMG amplitude envelopes across the gait cycle for right soleus (SOL_R_), right tibialis anterior (TA_R_), left soleus (SOL_L_), and left tibialis anterior (TA_L_) muscles. (A) Forward walking: comparison between Baseline (black line) and Washout (gray line) conditions, with shaded areas representing (*±*) one standard error of the mean. Yellow-highlighted region in SOL_R_ indicates a statistically significant cluster (*p* = 0.027) during early stance phase. (B) Bidirectional walking: EMG envelopes across blocks 1–4 (cyan, blue, magenta, and purple lines, respectively), with shaded areas representing (*±*) one standard error of the mean. Yellow-highlighted region in TA_L_ indicates a statistically significant cluster (*p* = 0.037) during late swing phase. Statistical comparisons were performed using cluster-based permutation tests with family-wise error rate correction.

During BDW, the right SOL showed a clear phase shift compared to baseline FW, with its activation peak occurring earlier in the gait cycle. This shifted pattern persisted across blocks 1-4 (colored lines) with activation magnitude gradually decreasing by block 4 (Figure 5 B, first panel). Right TA activation pattern shifted earlier in the gait cycle compared to baseline FW, with a marked increase in activation amplitude during block 1, followed by a gradual reduction through block 4 (Figure 5 B, second panel). Left SOL maintained similar timing as in its baseline FW across blocks 1-4, but early-stance activity increased in magnitude relative to baseline (Figure 5 B, third panel). Left TA maintained its bimodal structure throughout blocks 1-4, but the first (stance-related) burst was stronger and more prolonged relative to baseline (Figure 5 B, fourth panel). A statistically significant difference was observed only in the left TA during late swing phase (highlighted yellow region; p = 0.037), with no significant differences observed in the other muscles during BDW (Figure 5, Row B panels 1-3).

### 4.2 EMG burst duration relative to gait cycle

To examine whether BDW adaptation involved changes in the temporal structure of muscle activation (rhythm generation), we quantified burst duration relative to gait cycle duration for each conditions. The slope of the burst-to-cycle duration relationship varied across participants and conditions, with both positive and negative values observed, reflecting inter-individual variability in how muscle activation timing scaled with gait cycle duration (Figure 6). The ratio of burst duration to cycle duration was calculated for the SOL and TA muscles on both legs across the study, summarized in Figure 7.

**Figure 6:**
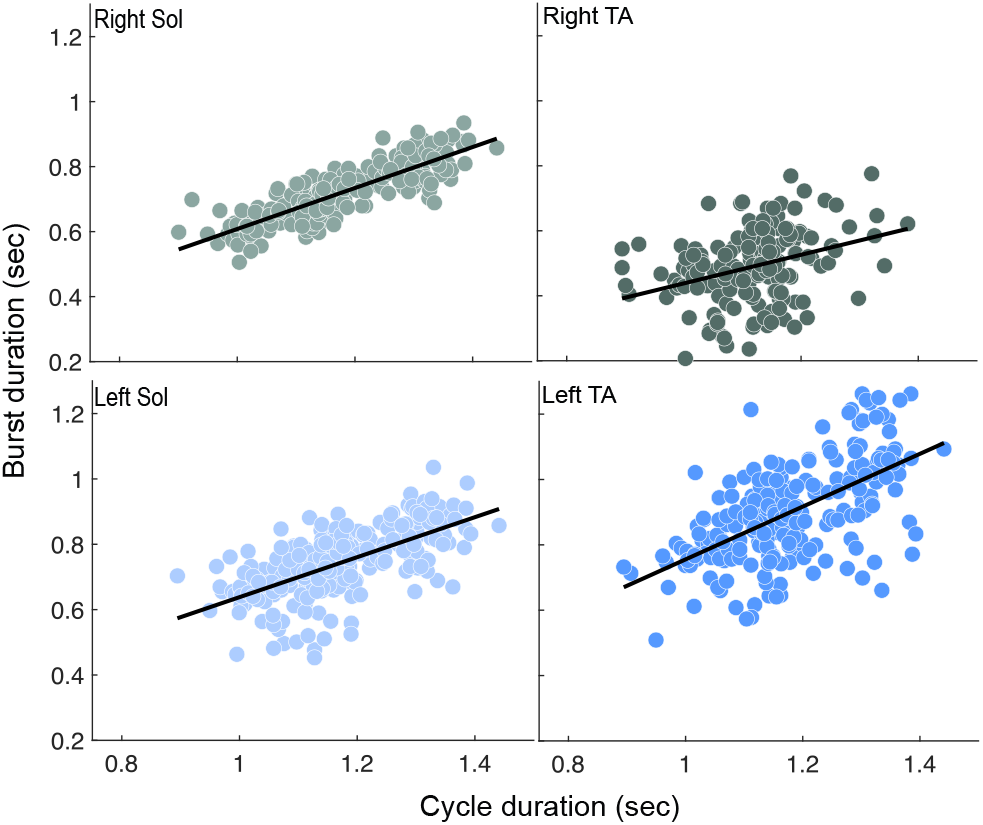
EMG burst detection to cycle duration. Relationship between burst duration and cycle duration for all four muscles during a one BDW block in a representative sample. Each point represents a single burst, and regression lines show positive associations between burst duration and cycle duration.

**Figure 7:**
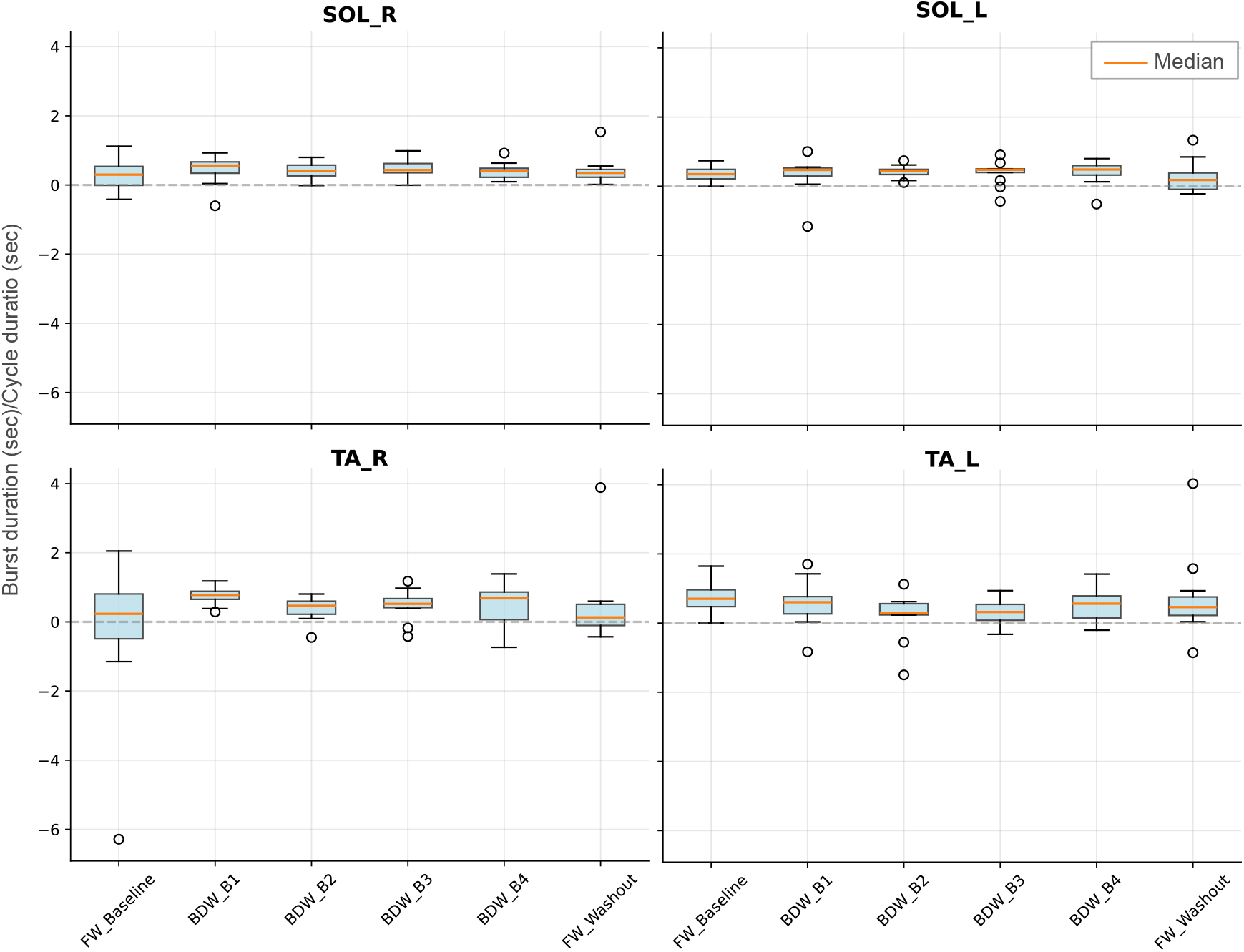
Burst-to-cycle duration ratio across forward and bidirectional walking conditions. Box plots show the ratio of burst duration (s) to cycle duration (s) for soleus and TA muscles bilaterally across the duration of the study. The orange line represents the median, boxes show the inter-quartile range (25th-75th percentile), whiskers extend to 1.5 times the inter-quartile range (IQR), and circles indicate outliers beyond this range.

The right SOL showed an increase in burst-to-cycle ratio from baseline FW to early BDW adaptation (block 1), which gradually decreased through blocks 2-4, and returned to near-baseline levels during washout (Figure 7, top left panel). The left SOL maintained relatively stable ratios throughout BDW blocks, with a decrease observed during the washout period (Figure 7, top right panel). The right TA showed an increase in burst-to-cycle ratio from baseline to early BDW (block 1), which remained elevated through block 4, and returned to near-baseline levels at washout (Figure 7, bottom left panel). The left TA exhibited a decrease in ratio from baseline to blocks 1 and 2, partial recovery by block 4, and intermediate values during washout (Figure 7, bottom right panel). None of these changes were statistically significant

#### Flexor-Extensor Dominance During Bidirectional Walking

To assess whether the two legs adopted different coordination strategies during BDW, we examined the relative dominance of flexor versus extensor muscle activation patterns by analyzing the relationship between EMG burst duration and cycle duration as described in [25] for the TA (ankle flexor) and SOL (ankle extensor). We classified trials as flexor-dominant when the ratio of TA burst-to-cycle duration to SOL burst-to-cycle duration exceeded 1, indicating relatively longer TA activation. Conversely, ratios less than 1 indicated extensor-dominant patterns. During the BDW blocks, the right leg (moving backward) and left leg (moving forward) showed generally similar patterns of flexor dominance. However, the left leg demonstrated notably more variable coordination in the third training block, with a higher occurrence of negative ratios compared to the right leg. At baseline, most trials displayed flexor-dominant patterns. During washout, there was a slight decrease in flexor dominance.

## 5 Discussion

### 5.1 Summary

Bidirectional walking (BDW), each leg moving at the same speed but in opposite directions, poses a unique challenge to the locomotor control system. This task requires maintaining anti-phasic inter-limb coordination while the two legs execute stance and swing in opposite directions. In our initial study [17], we characterized the spatiotemporal adaptations to BDW, including changes in step-length asymmetry, double-stance timing, and joint angles. In the present work we examined bilateral muscle activation profiles by quantifying temporal activation (timing/rhythm generation) and intensity (amplitude/pattern formation) of surface electromyography (EMG) signals from the soleus (SOL) and tibialis anterior (TA) muscles. Notably, the right SOL exhibited changes in burst-to-cycle duration ratio during early adaptation, while the right TA showed a corresponding phase shift and amplitude modulation both normalizing in later blocks and upon return to forward walking (FW). These results are consistent with and extend our earlier kinematic findings, demonstrating that humans dynamically recalibrate both muscle timing and amplitude to maintain coordinated locomotion under new environmental demands (e.g., BDW).

### 5.2 CPG Framework and Rhythm vs. Pattern Generation

The responsive flexibility required for BDW, or any other necessary locomotor adaptation, is managed by sophisticated neuromuscular coordination within the central nervous system (CNS); the tri-partite system of the spinal central pattern generator (CPG), brainstem, and cortical structures [1]. The CPG framework is functionally divided into a *rhythm generator*, responsible for the temporal structure and timing of muscle activation, and a *pattern formation* layer, which shapes the amplitude and intensity of muscle output by coordinating muscle pools [26, 27, 28, 29, 30]. Our study’s results strongly support this functional separation. The adaptation to BDW is achieved through modification of rhythm generator output, indicated by asymmetrical temporal adaptations in SOL and TA. The increase in burst-to-cycle duration ratio of the right SOL and TA during early adaptation reflects the neural strategy that restructures limb timing for reverse stepping. For example, the right SOL median burst-to-cycle ratio increased from 0.299 (95% CI: -0.019 to 0.621) to 0.560 (95% CI: 0.327 to 0.708) during early adaptation.

For pattern formation, the right SOL and TA showed clear phase shifts, with activation peaks occurring earlier during BDW. This change decreased gradually over blocks 1-4. Left SOL EMG maintained its timing but increased in strength just before stance and during early stance. The left TA kept its bimodal structure. Its pre-stance (i.e., late-swing) burst was larger.

These modulations show that locomotor CPG mechanisms are highly flexible and context-dependent [27, 28, 30]. The lack of statistical significance of the changes in rhythm generation metrics may be attributable to methodological constraints: fixed treadmill speed may have constrained temporal adjustments, and group averaging across 12 participants may have attenuated individual-level responses in addition to a relatively smaller sample size.

While adaptation relies on modulating the rhythm generation output, the core organizational structure of muscle synergies (the pattern-formation output) is often maintained across different locomotor challenges, such as changes in walking speed, loading, or even in the overall synergy components used for forward and backward walking in other studies [31, 32, 30, 33]. Further investigation of the pattern-formation layer requires muscle synergy analysis, which requires recording from a large number of muscles (e.g., studies tracking human adaptation in 15 muscles per leg or 13 leg muscles bilaterally [15, 34]. Recording only four muscles (SOL and TA bilaterally) restricted our ability to perform muscle synergy analysis to better describe changes in the pattern formation layer. We should note here that the present results are from the first session of a long-term study in which participants practiced BDW in three sessions/week for 8 weeks. Recording from more than four muscles (SOL and TA bilaterally) for so many sessions was not essential for the long-term study and would have constituted an excessive burden for the participants.

Ultimately, while the CPG provides the spinal foundation for rhythm and pattern formation, the expression and refinement of this complex BDW pattern in humans relies heavily on integration with descending inputs from the brainstem and cortical structures (including midbrain locomotor region (MLR) activation of reticulospinal pathways). These descending inputs modulate the CPG to provide the necessary directional drive and predictive control [35, 5, 12, 27].

### 5.3 Muscle Adaptation, Temporal Dynamics, and Cross-Species Comparison

FW and backward walking (BW) are remarkably similar motor behaviors (i.e., close to time-reversed mirror images). However, studies show mixed findings as to whether they share neural circuits [36, 37, 38, 39]. This independence implies a two-level control structure where the high-level networks responsible for adaptation and for encoding the timing of muscle activation (rhythm generation) are direction-specific, even if the lower-level muscle synergy modules (pattern formation) themselves are shared by FW and BW [32].

The asymmetric muscular demands of BDW initiated task-specific adaptations in both limbs, following a characteristic temporal progression similar to error-driven motor learning. Unlike typical split-belt paradigms where both legs move in the same direction at different speeds, BDW requires each leg to simultaneously execute fundamentally different movement strategies, one stepping forward and one stepping backward, making direct comparison with split-belt adaptation challenging. In the backward-moving right leg, the substantial but transient increase in the SOL burst-to-cycle duration ratio during early BDW Block 1 may serve as a critical neural strategy to generate force and prolong the stance phase so as to meet the new biomechanical demands of reverse stepping. Concurrently, the forward-moving left leg displayed a compensatory pattern, maintaining relatively stable timing in the SOL but showing stronger TA stance bursts and increased SOL early-stance activity (amplitude modulation). Analysis of flexor-extensor coordination indicates, while both legs maintained flexor dominance, the left leg showed significantly more variable coordination. This variability is consistent with the possibility that the left leg engaged in exploratory adjustments during mid-adaptation (BDW Blocks 2 & 3), actively searching for the optimal temporal coordination needed to synchronize with the constantly perturbed right limb. This adaptation followed a clear temporal progression, starting with the largest, immediate changes (Block 1: exploration/initial response) that progressively stabilized (BDW Blocks 2–3) until reaching a steady-state (Block 4: refinement/late adaptation). This time course is similar to the rapid onset and refinement seen in error-driven human split-belt learning [5, 15, 17, 33]. However, the persistence of kinematic aftereffects (e.g., negative step-length asymmetry) upon return to FW, contrasted with the finding that most EMG parameters, including the right SOL ratio, returned remarkably close to baseline during washout [17]. While we cannot determine the precise mechanisms from the present data, one plausible explanation is that the kinematic aftereffects may reflect adaptive adjustments in supraspinal feedback mechanisms. In contrast, the adaptations in spinal muscle timing patterns (the rhythm generator) disappear rapidly when the unique directional constraints are removed [15, 40].

EMG analysis in decerebrated cats show that hindlimb flexor and extensor muscles maintain reciprocal activation patterns during BDW [13]. A distinct asymmetry was observed in the forward-stepping limb: it took longer steps with greater joint angle ranges than the backward-stepping limb. In contrast, [12] found in BDW that the forward-stepping limb showed lower EMG amplitude in ankle extensor, hip flexor and knee extensor muscles, and lower extensor activity compared to either hindlimb during FW. The backward-stepping limb exhibited greater semitendinosus (hip extensor and knee flexor) EMG amplitude during BDW than during FW, but lower amplitude than during bilateral backward walking. These findings in cats provide a foundation for interpreting our human EMG results.

Animal studies show that spinal cats can perform BDW while maintaining coordination. This performance relies on a common rhythm-generating mechanism that preserves cycle durations when stepping direction changes [12]. However, cats (unlike humans) do not show locomotor adaptation: they do not return to symmetry with prolonged split-belt exposure and they show no aftereffects [29, 41]. This difference underscores the probable human reliance on supraspinal structures (such as the cerebellum) for refinement, preservation, and expression of BDW adaptation, especially for goal-directed tasks like maintaining stability during bipedal locomotion. In contrast, lower-level reciprocal coordination and cycle duration are maintained by the same spinal circuits [12].

This flexibility in BDW is also evident in early human development, further supporting the notion that the capacity for BDW is fundamental to the human locomotor system. Yang et al. [42] reported that 6 of 10 infants could perform BDW on a split-belt treadmill, suggesting remarkable flexibility in human neural control. This early-life capability suggests an autonomous pattern generator exists for each leg, with flexible coupling between the spinal rhythm and pattern generators for coordinating right and left legs [42]. While performing BDW, the infants maintained a consistent reciprocal relationship between their limbs; this ensured that the swing phases never overlapped [42]. These findings in cats and infants help in interpreting our human adult EMG results. They suggest that the capacity for BDW is always present at the spinal level. In adult humans, it is refined and adapted by supraspinal contributions.

### 5.4 Neuromuscular Basis of Kinematic Adaptations

Muscle activity adjustments underlie the kinematic changes previously reported [17], including bilateral step length reduction and inter-limb phasing shifts. For the backward-moving right leg, the ankle extensor (SOL) significantly increased its activation duration, apparently reflecting a command from the rhythm generator to prolong the stance phase. This temporal adjustment produced the hip, knee, and ankle joint angles needed for stable reverse stepping and contributed to reducing step length. Concurrently, the forward-moving left leg showed a compensatory pattern, including increased SOL early-stance activity. This modulation stabilized balance and regulated the asymmetric double stance duration (DS) (Figure 5 from [17]) which changed during BDW). This relationship indicates that the essential kinematic adaptations reflect changes in muscle timing that are driven by neural commands.

### 5.5 Limitations

First, we had 12 healthy participants. This modest number, combined with the high variability across participants may account for the lack of statistically significant changes in rhythm generation metrics. Second, we did not record from hip flexors and extensors. Third, like almost all previous studies of BDW, the analysis focuses on the data from a single session of BDW. Thus, it does not indicate the long-term impact of continued BDW on BDW itself, as well as on FW. As noted above, this session initiated a long-term multi-session study of BDW designed to described that long-term impact, and thereby address this limitation.

### 5.6 Conclusion

We studied the neuromuscular mechanisms underlying bidirectional walking by examining bilateral muscle activation patterns in the SOL and TA during BDW adaptation. We found that participants modulated both the temporal structure (rhythm generation) and amplitude characteristics (pattern formation) of muscle activation to accommodate the asymmetric demands of stepping in opposite directions. Specifically, the backward-moving leg showed increases in SOL burst-to-cycle duration ratios and TA phase shifts during early adaptation, while the forward-moving leg showed compensatory amplitude modulations. These muscle activation adjustments drove the spatiotemporal kinematic changes previously observed [17], including bilateral step-length reduction, altered inter-limb phasing, and asymmetric double stance timing. Analyses now underway of multi-session BDW will assess its long-term functional impact.

## Additional Information

### Data Availability

Data supporting the results of the study is available upon reasonable request.

### Author Contributions

All experiments were performed at the Samuel S. Stratton VA Medical Center, Albany, NY. The authors Helia Motjabavi (HM), Atra Ajdari (AA), Sebastian Rueda-Parra (SR), Darren E. Gemoets (DG), Jonathan R. Wolpaw (JW), and Russell L. Hardesty (RH) approve the accuracy and integrity of the study. The contributions of the each author are listed below.

- Conceptualization/Experimental Design: HM, RH, JW
- Data Collection: HM, RH
- Data Analysis: HM, RH, AA, SR, DG
- Writing/Manuscript Preparation: HM, RH, DG, AA, SR, JW

## Acknowledgements

The authors would like to acknowledge Dr. Jonathan S. Carp for his valuable scientific input. We would also like to acknowledge the National Center for Adaptive Neurotechnologies (NCAN) and the Samuel S. Stratton VA Medical Center for administrative, logistical support and use of their facilities. Drs. Wolpaw and Gemoets are employees of the Samuel S. Stratton VA Medical Center. The contents of this manuscript do not represent the views of the US Department of Veterans Affairs or the United States Government.

